# Inhibiting disulphide bonding in truncated tau297-391 results in enhanced self-assembly of tau into seed-competent assemblies

**DOI:** 10.1101/2025.02.05.636249

**Authors:** Sebastian S. Oakley, Karen E. Marshall, Georg Meisl, Alice Copsey, Mahmoud B. Maina, Robert Milton, Thomas Vorley, John M. D. Storey, Charles R. Harrington, Claude Wischik, Wei-Feng Xue, Louise C. Serpell

## Abstract

Tau undergoes fibrillogenesis in a group of neurodegenerative diseases termed tauopathies. Each tauopathy is characterized by tau fibrils with disease-specific conformations, highlighting the complexity of tau self-assembly. This has led to debate surrounding the precise mechanisms that govern the self-assembly of tau in disease, especially the involvement of disulphide bonding (DSB) between cysteine residues. In this study, we use a truncated form of tau, dGAE, capable of forming filaments identical to those in disease. We reveal the impact of DSB in dGAE assembly and propagation by resolving the global mechanisms that dominate its assembly. We found evidence for surface-mediated secondary nucleation and fragmentation being active in dGAE assembly. The inhibition of DSB during dGAE assembly leads to an enhanced aggregation rate through a reduced lag phase, but with no effect on the global assembly mechanisms. We suggest this is due to the formation a dominant, seed-competent species in the absence of DSB that facilitates elongation and secondary nucleation resulting in enhanced assembly. *In vitro* seeding assays reveal the recruitment of endogenous tau in a cell model only when using dGAE species formed under conditions that inhibit DSB. Our results further support the use of the *in vitro* dGAE tau aggregation model for investigating the mechanism of tau assembly, the effect of varying conditions on tau assembly and how these conditions affect the resultant species. Further studies may utilise dGAE and its aggregates to investigate tau seeding, propagation and to highlight or test potential targets for therapies that reduce the spread of pathologic tau throughout the brain.

## INTRODUCTION

Tauopathy is a collective term for a group of neurodegenerative diseases that are characterised by the deposition of abnormal tau aggregates throughout the brain. These diseases include Alzheimer’s disease (AD), frontal temporal dementia (FTD), chronic traumatic encephalopathy (CTE) and corticobasal degeneration (CBD) ^1^. Tau is a microtubule-associated protein involved in promoting and stabilising the microtubule network ^2^. However, in the brains of patients with tauopathies, tau undergoes a pathological self-assembly process to form highly ordered amyloid fibrils ^3^. These tau amyloid fibrils are strongly associated with neurodegeneration, cognitive decline and clinical dementia ^4^. Recent cryo-electron microscopy (cryo-EM) studies have resolved the atomic structure of the tau filament structures associated with different tauopathies ^5, 6, 7^ revealing that, in each disorder, tau self-assembles to form disease-specific conformations. Despite this, all filament structures resolved to date show that a similar region of tau associates to form the cross-beta core and that this is comprised predominantly of the imperfect repeat region ^8^. This illustrates the complex polymorphic capacity of tau to form multiple specific atomic structures and interactions, which are associated with disease-specific and clinically distinct cognitive impairments. Revealing the initial stages and the mechanisms involved in tau self-assembly, and how they affect the final fibril structure, is crucial for our understanding of tauopathies and in the development of therapeutics.

Amyloid-specific dyes, such as Thioflavin T or S (ThT/ThS), have been used in kinetic assays to analyse the rate of protein self-assembly into filaments for a range of amyloidogenic proteins, such as β_2_-microglobulin (β_2_m) ^9^, tau ^10^, α-synuclein ^11^ and amyloid-β ^12^. The traces obtained from ThT/ThS kinetic assays of these proteins have been pivotal in revealing the mechanisms that dominate the process of amyloidogenic protein self-assembly, such as primary or secondary nucleation processes ^9, 13, 14^. Primary nucleation refers to a monomer-only nucleation process, whereby the assembly of small oligomeric aggregates, or nuclei, depends on the rates of association of monomers and its dissociation back to monomers. Secondary nucleation denotes a fibril surface-dependent nucleation process, where the assembly of nuclei from monomers is catalysed by pre-existing fibrils formed from the same protein monomer ^13, 15^. Fragmentation is a form of secondary process that amplifies fibril formation through repeated division and elongation cycles. Existing aggregates fragment, exposing additional fibril ends that facilitate further elongation ^9, 16^.

ThS/T aggregation kinetic assays can also be utilised to explore the role of specific bonds, bonding regions, and residues in amyloid assembly. The impact of disulphide bonding (DSB) between cysteine residues for tau self-assembly is still highly debated. With the use of *in vitro* tau aggregation models, such as full-length tau (T40) and K18/K19 fragments templated using heparin, DSB was proposed as an essential step in tau self-assembly and propagation ^17^. However, the ability of these models to produce reliable tau filaments with the same macromolecular structure as those isolated from tauopathies has been disputed by cryo-EM studies ^18^. A truncated form of tau protein corresponding to residues 297-391, termed dGAE, assembles readily in reducing conditions ^19, 20^ to form twisted filaments that are structurally identical to AD paired helical filaments (PHFs) and CTE type II filaments ^21–23^. Under reducing conditions, cys322 is prevented from forming disulphide bonds and the variant C322A is useful in determining the contribution of the cystine residue. dGAE is therefore a valuable model for the investigation of tau self-assembly, filament structure and propagation.

In this study, we utilise ThS kinetics assays to investigate the assembly mechanisms of dGAE and dGAE-C322A, the involvement of DSB in dGAE assembly and the seeding characteristics of dGAE aggregates in a cellular environment. We show that secondary processes dominate the assembly mechanism of dGAE, with evidence showing that complex surface-mediated secondary nucleation and fragmentation are active in dGAE assembly. In addition, the inhibition of DSB in dGAE aggregation has no effect on the overall global assembly mechanism, but significantly enhances the rate of aggregation resulting in a shorter lag phase. Self-assembled species without DSB are significantly better at accelerating the aggregation process *in vitro* and at recruiting endogenous tau in FRET Biosensor cells. This suggests that the presence of DSB in dGAE filaments is detrimental for those aggregates to facilitate the propagation of tau fibril formation, and that the inhibition of DSB as an important step in the formation of pathological tau aggregates.

## MATERIALS AND METHODS

### Purification of truncated tau 297-391 (dGAE)

dGAE and dGAE-C322A proteins were expressed in *E. coli* BL21 cells grown in 2x Yeast Tryptone (2xYT) media supplemented with ampicillin. Over-night cultures were diluted to an OD600 of 0.01 and grown at 37°C, 250 rpm until they reached an OD600 of 0.6. at which point protein expression was induced by the addition of Isopropyl β-D-1-thiogalactopyranoside (IPTG) to a concentration of 0.4mM. After 2 hours induction, cells were harvested by centrifugation at 7,000 rpm for 10 minutes at 4°C (Beckman Avanti J-30I, JLA-16.25 rotor). Cell pellets were transferred to 50ml falcon tubes in 0.9% NaCl and cells were collected by centrifugation at 4,500 rpm for 10 minutes at 4°C (Beckman Avanti J-30I, JA 25.50 rotor). Cells from 4 litres of expression culture were resuspended in 80mls lysis buffer (50 mM MES, pH 6.25) with 1 mM Ethylenediaminetetraacetic acid (EDTA), 1mM dithiothreitol (DTT) supplemented with EDTA-free protease inhibitors (Roche). Cells were lysed using 3 min sonication (5s on/5s off) at 50% amplitude with 13-mm probe on ice and centrifuged at 10,000 rpm (Thermo Scientific Heraeus Multifuge 3S-R) to remove intact cells and cell debris. NaCl was added to the supernatant at 1.75g per 40ml plus DTT to a final concentration of 1mM. The cell suspension was boiled for 5 min to precipitate proteins which were removed by centrifugation at 10,000 rpm for 10 min at 4°C. The dGAE containing supernatant was dialysed over-night into MES buffer (50 mM MES (pH 6.25), 1mM ethylene glycol-bis(2-aminoethylether)-N,N,N’,N’-tetraacetic acid (EGTA), 5 mM EDTA, 0.2 mM MgCl_2_ and 5mM mercaptoethanol). dGAE was further purified by passage through a 5 ml HiTrap SP Sepharose column attached to an ÄKTA Fast Protein Liquid Chromatography (FPLC). The column was equilibrated at a flow rate of 1 ml min^−1^ in MES buffer. Protein was bound to the column at a flow rate of 0.5 ml min^−1^ and washed at a flow rate of 1 ml min^−1^ until the absorbance at 280 nm had returned to the baseline. Protein was eluted with a 10-times the column volume of MES buffer supplemented with 1M KCl and peak fractions were collected in a 96 deep-well plate. Fractions containing dGAE were pooled, dialysed overnight into phosphate buffer and stored at -80°C. The protein was then diluted in phosphate buffer (PB) (10mM; pH 7.4) for further experimentation.

### ThS kinetics and analysis

dGAE monomer was added at varying concentrations (1-20 µM) in degassed phosphate buffer, pH7.4 with 20µM Thioflavin S (ThS) in low-bind tubes. ThS is an analogue of ThT and was used instead of ThT because of its higher sensitivity with tau amyloid aggregates. Each sample was transferred to a non-binding µClear® bottom, black 96-well plate (Greiner Bio-one, 655906) with a foil seal to stop evaporation. The plate was incubated in a Molecular Devices SpectraMax i3x plate reader equilibrated at 37°C with high orbital shaking (469 rpm) and readings taken from the bottom of the plate every 5 minutes using an excitation filter at 440nm and an emission filter 483nm, to monitor ThS fluorescence intensity. The data were normalised using the initial plateau and final plateau using MATLAB as previously described ^9^. Normalised data were then subjected to global chemical kinetics analysis using Amylofit software ^14^ to observe how the data fit with each global model of aggregation.

### Preparation of dGAE fibrils

dGAE or dGAE-C322A (400µM) was diluted in 10mM PB +/- 10mM DTT ^21, 23^ and incubated at 37°C whilst agitating at a speed of 400rpm on an Eppendorf ThermoMixer® for 4d. The samples were then centrifuged at 16,000 g at 4°C for 30min. The supernatant was removed to provide protein concentration of the supernatant to estimate the protein content in the pellet. Protein concentration was estimated using the Bicinchoninic acid (BCA) assay. PB (10mM) was added to the pellet to suspend fibrils at a final concentration of 400µM dGAE. If the samples had DTT in the assembly mixture, these fibrils were washed once with PB, with an additional centrifugation step before final suspension. Sonication was done in a water bath sonicator for 10m with ice to reduce the effect of heating.

### Transmission electron microscopy (TEM)

Electron microscopy grids were prepared by placing 4 μl of sample onto Formvar®/carbon-coated 400-mesh copper grids (Agar Scientific), blotting excess and then washing with 4 μl of 0.22-μM filtered milli-Q water. Uranyl acetate (4 μl of 2% in water) was placed on the grid once for 1 minute and then blotted and the grid allowed to air-dry. TEM projection images were collected using a JEOL JEM1400-Plus Transmission Electron Microscope operated at 120 kV equipped with a Gatan OneView camera (4k × 4k). Images were recorded at 25 fps with drift correction using GMS3.

### Circular dichroism (CD)

dGAE and dGAE C322A fibrils were spun to separate the supernatant and pellet. The pellet was resuspended in PB to a final concentration of 400µM. CD was performed using a Jasco Spectrometer J715 and spectra collected in triplicate at a maintained temperature of 21 °C. Protein samples of the pellet (60µl) were placed into 0.2-mm path length quartz cuvettes (Hellma) and data were collected between wavelengths of 180 to 350 nm.

### Proteinase K (PK) digestion

Fibrils from each condition were diluted to 200µM with 10mM PB containing 25µg/ml of proteinase K (PK) (prepared in 50mM Tris/1mM CaCl_2_, pH7.5) and incubated for 1h at 37°C. The same amount of PK buffer was added to controls without PK. Samples were then subjected to SDS-PAGE electrophoresis. Samples were loaded onto an Any kDa Mini-PROTEAN® precast gel (Bio-Rad Laboratories) and run at 200V for 30min in TRIS-glycine-SDS running buffer. Coomassie stain was applied for 1h, destained once for 1h and again overnight. Gels were imaged on a Li-Cor Odyssey FC imaging system (exposure 30s).

### Cell culture and FRET aggregation assay

Tau RD P301S FRET Biosensor from ATCC (CRL-3275™) ^24^ were grown in Dulbecco’s modified Eagle medium/Nutrient Mixture F-12 (DMEM/F-12) supplemented with 10% foetal calf serum, 1% penicillin/streptomycin (P/S) and 1% L-glutamine. Cells were plated at a density of 15,000 cells per well in a 96-well plate and incubated for 2 days. 10µM of soluble, fibrils or sonicated fibrils from each condition (dGAE, dGAE+DTT, and dGAE-C322A) were added to the cells in growth media and incubated for 3d at 37°C in 5% CO_2_. Post 72h incubation, the plate was imaged using the automated Molecular Devices ImageXpress Pico using FITC filter with 20x objective. Analysis was carried out using the CellReporterXpress software, whereby optimisation of the cell counting analysis protocol was adapted to specifically select the punctate FRET signal produced as a result of aggregated endogenous tau. Parameters used were as follows: minimum size = 2, maximum size = 10, intensity = 150. This was optimised to be effective at isolating the FRET signal for accurate analysis of aggregated endogenous tau (Figure 1).

**Figure 1.**
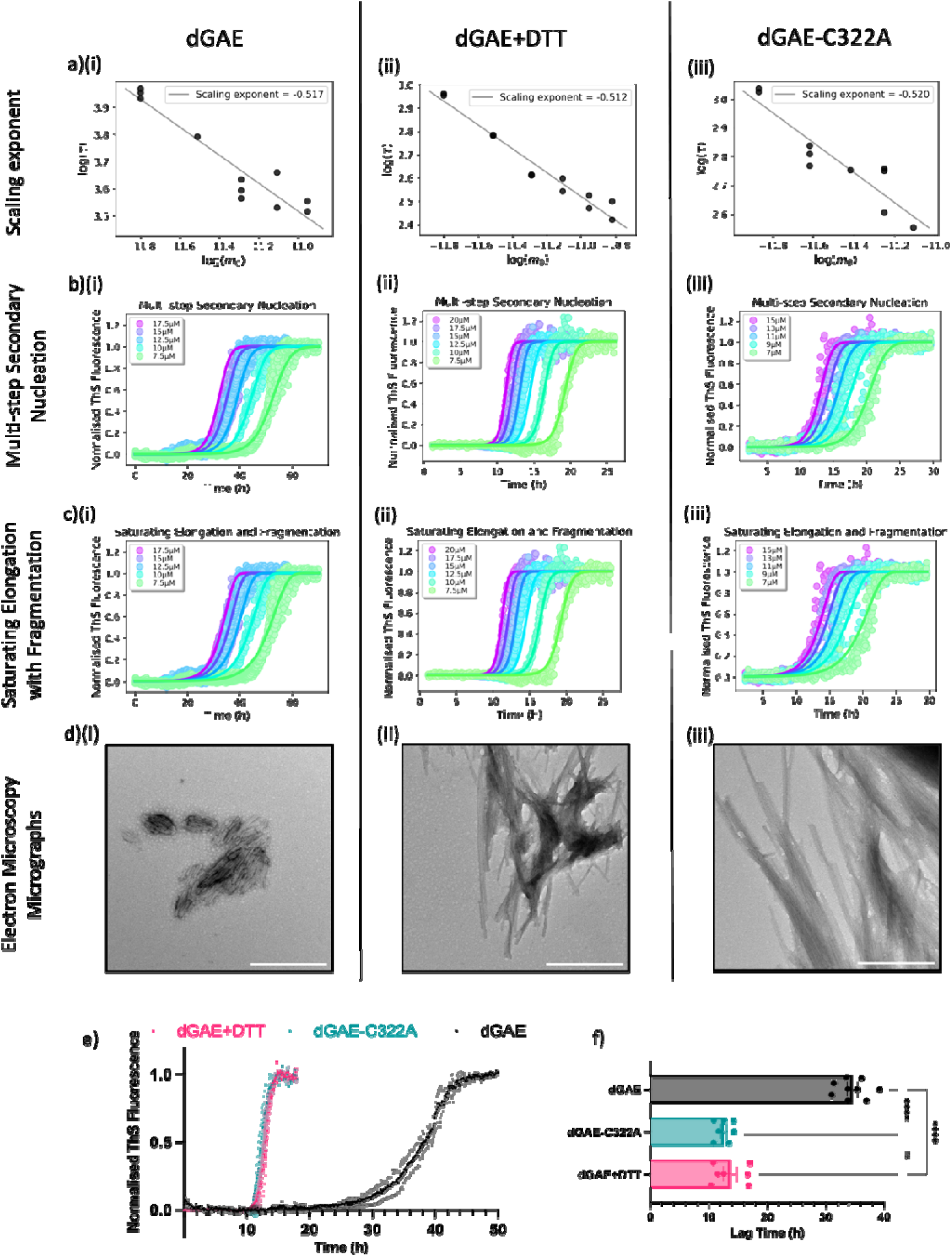
Impact of DSB on dGAE assembly. dGAE assembled in non-reducing conditions is shown in the left panels (i), dGAE assembled in reducing conditions with 10mM DTT in middle panels (ii), and dGAE-C322A assembled in non-reducing conditions in the right panels (iii). (a) Scaling exponent of each titration experiments. Normalised kinetic profiles plotted against different models of assembly: (b) multi-step secondary nucleation and (c) saturating elongation with fragmentation. (d i-iii) Electron micrographs of the resultant assemblies taken when the final plateau has been reached. Scale bars = 500nm. (e) Comparison of normalised thioflavin-S kinetic profile from one experiment with a starting concentration of 15µM monomeric dGAE in non-reducing conditions (black), dGAE in reducing conditions with 10mM DTT (pink), and dGAE-C322A in non-reducing conditions (green). (f) Quantification and comparison of the lag time for each condition; results were taken from 3 independent experiments. One-way ANOVA showed a significant difference between the groups (F =206.2, R2=0.9538, p <0.0001). Tukey’s multiple comparisons test showed significance when comparing dGAE (34.60h ± 0.8734h) with both dGAE+DTT (13.69h ± 1.139h, p < 0.0001) and dGAE-C322A (12.60h ± 0.5575h, p < 0.0001), but there was no significant difference between dGAE+DTT and dGAE-C322A.

### Data analysis and representation

Data and statistical analyses were performed using Microsoft Excel, GraphPad Prism 7 and MATLAB R2022a. All data are expressed as the mean ± SEM. When comparing two groups, a form of t-test was used to determine statistical significance. When comparing more than two groups, a form of one-way ANOVA test was used to determine if there is a difference between experimental groups and a control group. The normality and distribution of the data were calculated to decide upon the specific t-test and one-way ANOVA. Specific details for the individual statistical tests and multiple comparison tests performed can be found within the figure legends. Differences were considered to be statistically significant if *p* < 0.05.

## RESULTS AND DISCUSSION

### dGAE assembly is dominated by secondary processes and is independent of disulphide bonding

To investigate the contribution of DSB to dGAE assembly mechanisms, ThS fluorescence assays were used to monitor the kinetics of self-assembly of dGAE under conditions that allow or inhibit DSB. Conditions for reproducible spontaneous assembly of dGAE in non-reducing and reducing conditions (with 10mM DTT), or dGAE-C322A in non-reducing conditions were optimised using a SpectraMax i3x plate reader. dGAE-C322A variant was used to inhibit disulphide bonding through the substitution of the only cysteine residue in dGAE. This variant has previously been shown to assemble into filaments ^19^. dGAE samples prepared with a dilution series (1-20µM) were measured using ThS. Agitation was found to be necessary for reproducible assembly. All data showed sigmoidal-like appearance with a lag-phase, a steep growth phase, and a final plateau (Figure 1). A clear concentration dependence was observed showing faster assembly at higher concentrations, with unreliable assembly observed at a dGAE monomer concentration of ∼7µM (data not shown). The data were normalised to the initial baseline and final plateau before uploading to Amylofit software ^14^. The time to reach the midpoint between the initial baseline and final plateau values is known as the half-time which, when plotted against monomer concentration in a log-log plot, was used to generate the scaling exponent for dGAE assembly in the different conditions (Figure 1a). The straight line of the scaling exponent indicates that the dominant mechanism of aggregation does not change within the monomer concentration range tested. The value of the scaling exponent is close to -1/2 for all conditions (dGAE = -0.517, dGAE+DTT = -0.512 and dGAE-C322A = -0.520), which suggests that either or both fragmentation and saturated secondary nucleation (multi-step secondary nucleation) dominate dGAE aggregation, regardless of DSB involvement. To further clarify this, normalised kinetic profiles were plotted together with best-fit global models of aggregation mechanisms. A good fit of the model to the data was observed for both 1) multi-step secondary nucleation of monomers mediated by the fibril surface (saturated secondary nucleation) (Figure 1b) and 2) fragmentation process with primary nucleation and elongation mediated by the exposed ends of the fibrils (Figure 1c). Models for nucleation elongation, secondary nucleation, and saturating elongation and secondary nucleation show a poor fit for each condition (Sup. Figure 1).

The rate constants obtained from the global fitted models reveal a substantial increase in the rate of the secondary process when DSB formation is inhibited either through addition of DTT or with the dGAE-C322A mutation (Sup. Figure 2a and b), and this is predominantly responsible for the faster assembly observed when DSB is inhibited. The rate of primary nucleation is also affected, although in different ways. While the C322A mutation resulted in a slight increase in the rate of primary nucleation, addition of DTT leads to a significant decrease in the rate of primary nucleation (Sup. Figure 2c). Secondary processes dominate in all cases, so the effect of primary nucleation on the overall speed of the assembly is less pronounced (Sup. Figure 2d). However, the decreased importance of primary nucleation upon DTT addition is evident in the assembly curves becoming sharper, with a more sudden increase after a flat lag phase.

TEM was carried out to observe the morphology of fibrils formed under different conditions. dGAE in non-reducing conditions formed short fibrillar species prone to lateral association (Figure 1d)(i)). In contrast, a higher proportion of elongated fibrils were observed in the reducing conditions (Figure 1d)(ii)) or using dGAE-C322A (Figure 1d)(iii)) and this is consistent with previous work ^19^. A direct comparison of ThS fluorescence traces from 15µM of monomeric dGAE with or without DSB (Figure 1e), shows the significantly shorter in the lag phase of ∼11-12h when DSB is inhibited (dGAE+DTT or dGAE C322A) and ∼20h for non-reduced dGAE (Figure 1f).

These mechanistic investigations illustrate that the assembly of dGAE is governed by fibril-dependent secondary processes, as complex secondary nucleation processes on the surface of fibrils in solution through fragmentation or both. Although the overall mechanism is independent of DSB, the inhibition of these DSB leads to an enhanced assembly reaction, with a significant reduction in the lag-phase of assembly.

### dGAE species formed through the inhibition of DSB are more capable of seeding assembly of dGAE

Monitoring the assembly of proteins in the presence of preformed fibrils enables further examination of the involvement of secondary processes (secondary nucleation and fragmentation) by bypassing primary nucleation ^10^. Furthermore, it provides information on the differences in heterogenous seeding capabilities between assemblies formed with DSB (DSB(+)) in the dGAE sample, or prevention of DSB (DSB(-)) in the dGAE+DTT and dGAE-C322A samples.

The seeds were formed through previously optimised conditions and methods ^19, 21, 25^, and to mimic the conditions used in recent cryo-EM studies using dGAE ^23^. dGAE (400µM), with or without DTT (10mM), or dGAE-C322A (400µM) were agitated in an Eppendorf ThermoMixer® at 37°C for 4d. Assemblies were isolated, and their concentration estimated, as described in Materials and Methods. The fibrils were sonicated before being added at 1%, 5% and 10% of the initial monomeric protein concentration, 10µM. Following sonication, TEM images showed short fibrils with no observable differences in morphology between the conditions (Figure 2a). dGAE and dGAE+DTT seeds were added to wildtype dGAE monomer in a reducing environment (10mM DTT) because these conditions resulted in more reproducible ThS kinetic traces. dGAE-C322A seeds were added to dGAE-C322A monomer in non-reducing conditions.

**Figure 2:**
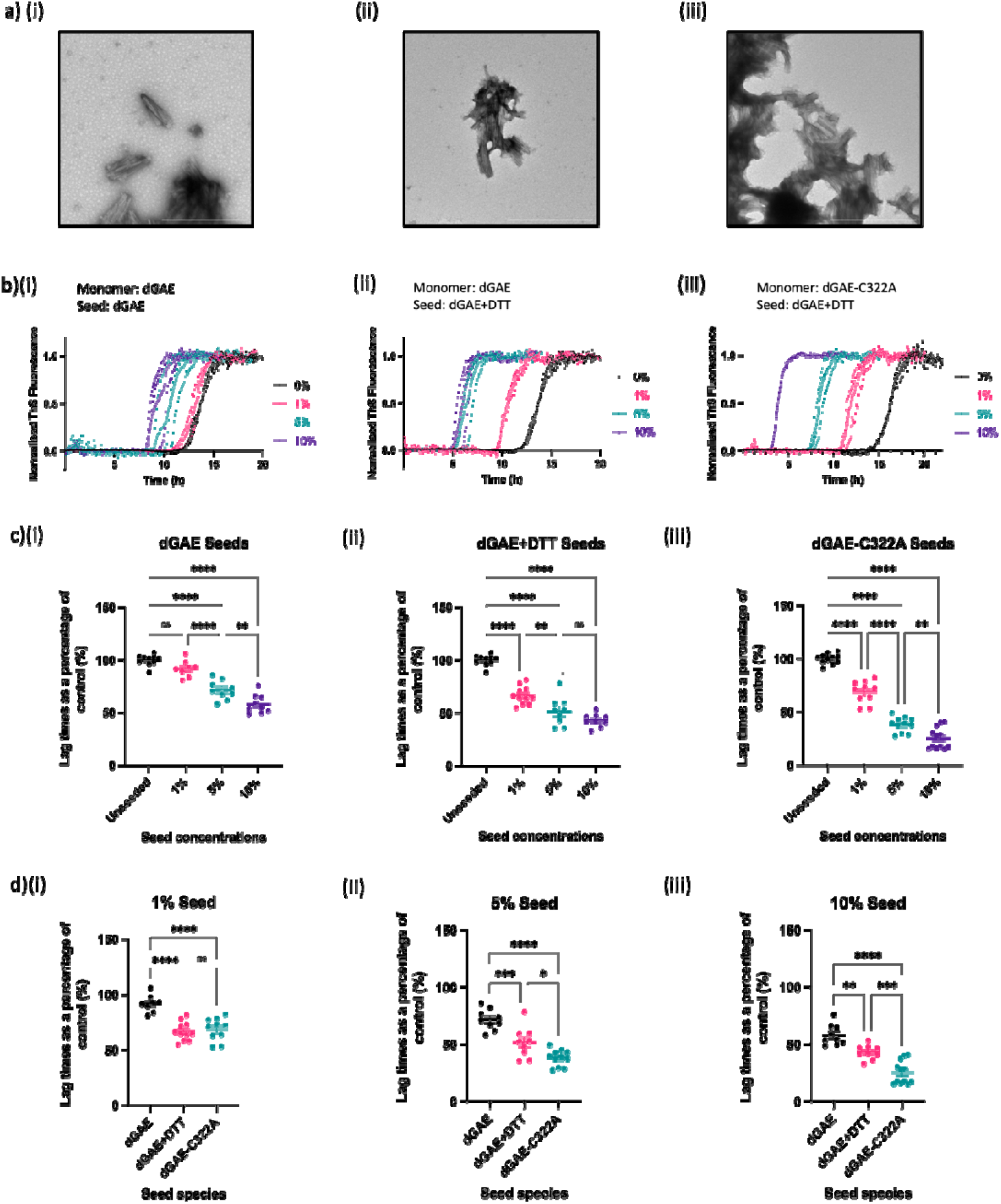
Non-disulphide species are more seed capable than disulphide species. Electron micrographs of seeds produced from sonicating fibrils produced in each condition, (a i) dGAE non-reducing conditions, (a ii) dGAE+DTT reducing conditions with 10mM DTT, and (a iii) dGAE-C322A non-reduced conditions. Scale bar represents 500nm. Examples of normalised thioflavin-S kinetics from a single experiment showing the seeding capability of each condition seed at 0% (control - black), 1% (pink), 5% (green) and 10% (purple) for dGAE seeds (b i), dGAE+DTT (b ii), dGAE-C322A (b iii). (c) Quantification and comparison of the lag times as a percentage of the unseeded control when 1%, 5% and 10% of seeds from each condition. (c i) Seeds produced from dGAE in non-reducing conditions added to 10µM of dGAE with 10mM DTT. One-way ANOVA shows significant difference between groups (F=54.41, R^2^=0.8361, p < 0.0001) from 4 independent tests. Tukey’s multiple comparison test shows 1% dGAE seed (92.07% ± 2.750%) does not induce a significant reduction in lag phase when compared to the control (100% ± 1.608%, p < 0.0001). Whereas the addition of 5% (71.77% ± 3.001%) and 10% (58.24% ± 3.023%) dGAE seeds does induce a significant reduction in lag phase when compared to the control (p < 0.0001). Significant reduction in the lag phase is also seen with the addition of 5% when compared to 1% dGAE seeds (p < 0.0001), and when comparing 10% to 5% dGAE seeds (p = 0.0047). (c ii) Seeds produced from dGAE in reducing conditions (10mM DTT) added to 10µM of dGAE with 10mM DTT. One-way ANOVA shows significant difference between groups (F=77.92, R^2^=0.8698, p < 0.0001) from 4 independent tests. Tukey’s multiple comparison test shows 1% (66.97% ± 2.557%), 5% (51.76% ± 4.479%) and 10% (43.20% ± 2.069%) dGAE+DTT seed does induce a significant reduction in lag phase when compared to the control (100% ± 1.608%, p < 0.0001). Significant reduction in the lag phase is also seen with the addition of 5% when compared to 1% dGAE seeds (p = 0.0026), but not between 10% and 5% dGAE seeds (p = 0.1847). (c iii) Seeds produced from dGAE-C322A in non-reducing conditions added to 10µM of dGAE-C322A. One-way ANOVA shows significant difference between groups (F=179.6, R^2^=0.9309, p < 0.0001) from 4 independent tests. Tukey’s multiple comparison test shows 1% (68.92% ± 3.247%), 5% (38.25% ± 2.127%) and 10% (25.47% ± 2.920%) dGAE+DTT seed does induce a significant reduction in lag phase when compared to the control (100% ± 1.399%, p < 0.0001). Significant reduction in the lag phase is also seen with the addition of 5% seed when compared to 1% dGAE seeds (p < 0.0001), and between 10% and 5% dGAE seeds (p = 0.0038). (d) Comparison of percentage lag times induced with the seeds produced from the different conditions at 1% (di), 5% (dii) and 10% (diii) using the same data from (c). (d i) One-way ANOVA shows significant difference between groups for 1% seed addition (F=36.20, R^2^=0.7283, p < 0.0001) from 4 independent tests. Tukey’s multiple comparison test shows a significant reduction in lag phase with the addition of dGAE+DTT and dGAE-C322A when compared to dGAE (p < 0.0001), but no significant difference observed between dGAE+DTT and dGAE-C322A seeds (p = 0.8734). (d ii) One-way ANOVA shows significant difference between groups for 5% seed addition (F=36.20, R^2^=0.7283, p < 0.0001) from 4 independent tests. Tukey’s multiple comparison test shows a significant reduction in lag phase with the addition of dGAE+DTT and dGAE-C322A when compared to dGAE (p < 0.0001), as well as significant difference observed between dGAE+DTT and dGAE-C322A seeds (p = 0.0153). (d iii) One-way ANOVA shows significant difference between groups for 10% seed addition (F=36.20, R^2^=0.7283, p < 0.0001) from 4 independent tests. Tukey’s multiple comparison test shows a significant reduction in lag phase with the addition of dGAE+DTT when compared to dGAE (p = 0.0033), a significant reduction with the addition of dGAE-C322A when compared to dGAE (p < 0.0001), as well as significant difference observed between dGAE+DTT and dGAE-C322A seeds (p = 0.0003).

The addition of 1% dGAE seeds to monomeric dGAE resulted in no significant reduction in the lag phase, whereas the addition of 5% and 10% seeds results in a significant reduction when compared to the unseeded control (Figure 2b i and c i). A significant reduction in lag phase was observed when dGAE+DTT seeds (Figure 2b ii and c ii) or dGAE-C322A seeds (Figure 2b iii and c iii) were added at all percentages. These data show a clear reduction in the aggregation lag phase with the addition of preformed aggregates, providing clear evidence for the involvement of secondary nucleation in dGAE and dGAE-C322A assembly. The reduction in lag phase is generally significantly concentration dependent (Figure 2b and c). Although the reduction in lag phase for the increase from 5% to 10% dGAE+DTT was not significant.

Further analysis compared the normalised data as a percentage of the unseeded control (Figure 2d). The addition of 1% seed dGAE+DTT and dGAE-C322A seeds resulted in a significant reduction in lag-phase when compared the addition of dGAE seeds (Figure2d i). 5% and 10% seeds resulted in significant difference between the three groups whereby dGAE-C322A seeds significantly shorten the lag phase the most and dGAE-DTT the least (Figure 2d ii and 2d iii). To summarise, these data showed that DSB(-) dGAE species (dGAE+DTT and dGAE-C322A seeds) were significantly more effective at seeding *in vitro* dGAE assembly when compared to DSB(+) dGAE species, with dGAE-C322A seeds as the most effective seed.

Furthermore, the data also demonstrate that the introduction of sonicated, fibrillar dGAE species induced a reduction in the assembly lag phase, but it did not eliminate the lag phase completely, even at 10% of monomeric concentration. This is consistent with that view that surface-mediated nucleation dominates the seeding reactions ^26^, which supports the involvement of multi-step secondary nucleation (Figure 1b). Additional seeding experiments showed that dGAE fibrils after sonication to produce smaller truncated fibrils, are more competent seeds when compared to long mature fibrils (Sup. Figure 3). This could be evidence for the involvement of fragmentation in dGAE assembly, because of the increased presence of exposed fibril ends facilitating the repeated cycles of elongation and fragmentation. However, this could be due to sonication helping to reduce clumping in the sample that facilitates the more competent seeding observed in sonicated samples, and not the length of the fibrils being the important factor.

The reduction in lag phase with the addition of preformed seeds confirms the presence of secondary processes in the assembly of dGAE at all conditions. More importantly, there was a difference in the ability of the seeds to accelerate the assembly of dGAE, when comparing seeds formed in conditions favouring DSB and with those preventing DSB. Our results suggest that DSB(-) dGAE seeds are more capable of seeding the *in vitro* assembly of dGAE.

### The disulphide bond influences the macromolecular structure of the dGAE filaments

Cryo-EM revealing the structurally distinct tau filaments seen in tauopathies have highlighted the relationship between amyloid filament polymorphic structures and disease ^5, 6, 8^. Next, we focussed on investigating the differences in structure of dGAE filaments formed in conditions favouring and inhibiting DSB to gain insights into the effect of conditions on polymorphs.

Fibrils were assembled using 400µM of monomeric protein, as described in materials and methods. Lower starting monomer concentrations showed that prevention of DSB resulted in longer fibrils (Figure 1d ii and iii). At 400µM, electron micrographs show that reducing conditions result in filaments that are longer than those formed in non-reducing conditions (Figure 3a i and ii). The reducing conditions result in a mixture of twisted filaments that mimic PHF (red arrows) ^21, 23^, and straight filaments that laterally associate to form thicker filaments (blue arrows). dGAE-C322A monomer assembles to form straight laterally associated filaments, that resemble the filaments seen in the dGAE+DTT sample shown with the blue arrows, but PHF-like twisted filaments were not observed (Figure 3a iii). A higher starting monomer concentration (400µM) resulted in samples that contained mostly elongated filaments (Figure 3a i-iii), whereas filaments formed using low concentrations (10µM) withdrawn from kinetics assays were overall shorter and contained less elongated fibrils (Figure 1d i-iii).

**Figure 3:**
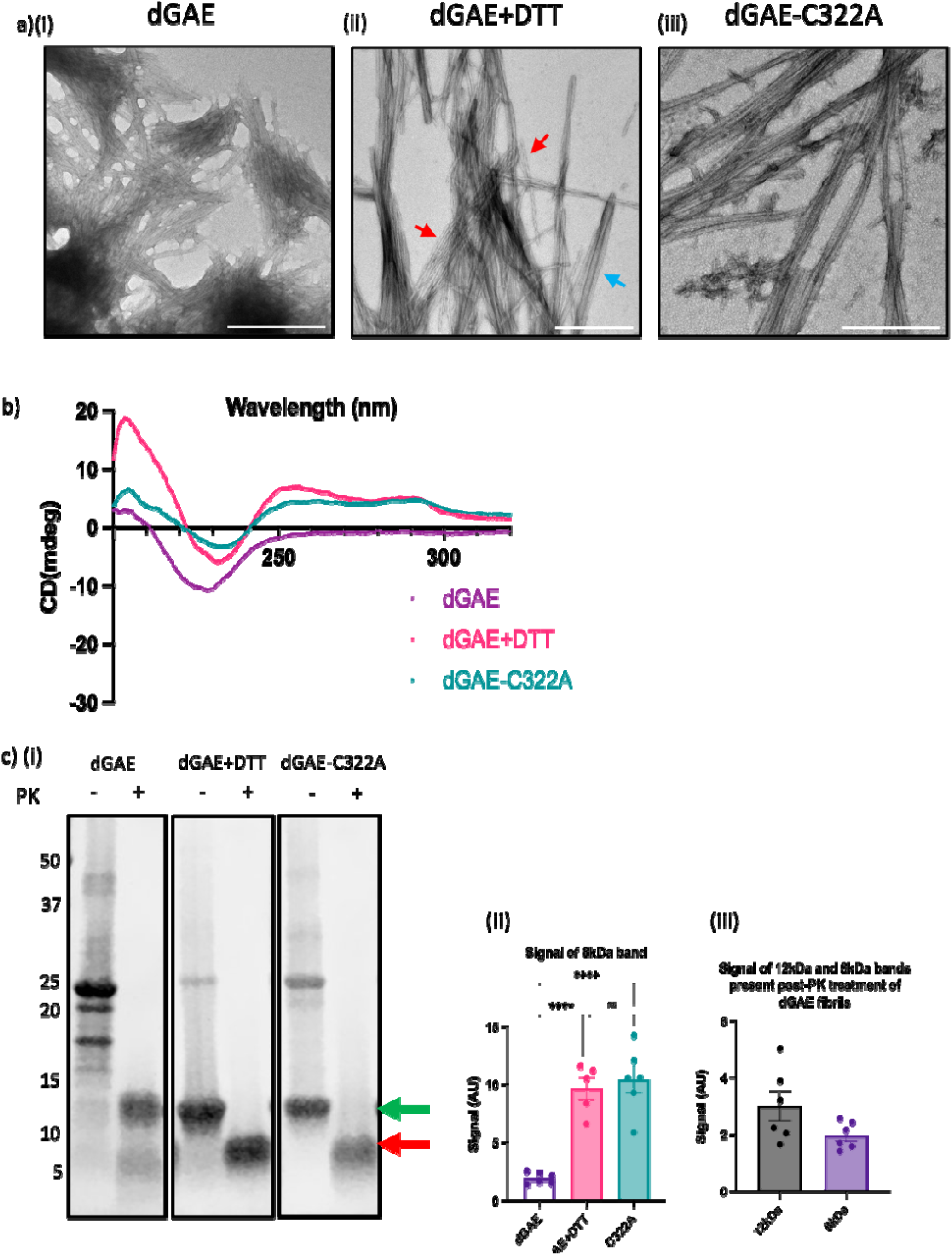
Macromolecular differences between fibrils assembled in conditions that favour disulphide bonding compared to fibrils assembled in conditions inhibiting disulphide bonding Electron micrographs of fibrils formed from 400μM dGAE in non-reducing conditions (a i) and in reducing conditions with 10mM DTT (a ii) and 400μM dGAE-C322A in non-reducing conditions (a iii). Scale bar represents 500nm. (b) CD spectrum from the pellet from each condition. (c i) SDS-PAGE and Coomassie staining for 9μl of 200μM fibrils from each condition with and without proteinase K treatment. Arrows represent the difference between the 12-kDa (green) and 8-kDa (red) monomers. (c ii) Quantification and comparison of the signal from the 8-kDa band for each condition after proteinase K treatment. One-way ANOVA showed significant difference between each group (F=30.64, R^2^=0.8140, p < 0.0001) from 5 independent tests. Holm-Šídák’s multiple comparisons test showed significant increase in the signal from the 8-kDa band in the dGAE+DTT sample (9.686 ± 0.9527) and dGAE-C332A sample (10.45 ± 1.150) when compared to the dGAE sample (1.973 ± 0.1982, p < 0.0001). There is no significant difference between dGAE+DTT and dGAE-C322A. (c iii) Quantification and comparison of the signal from the 12-kDa and 8-kDa band from the dGAE sample. Unpaired t test showed no significant difference between each group (F=7.072, R^2^=0.2516, p = 0.0966) from 6 independent tests.

The CD spectra of dGAE filaments after the fibrils have been isolated through centrifugation, showed a minimum at ∼225nm and a maximum at ∼200nm, arising from a high β-sheet content as expected for fibrils (Figure 3b) ^19^. The DSB(-) fibrils also show this strong β-sheet signal, but with two additional maxima at ∼250nm and ∼290nm, which suggests an increased order for these filaments compared to the dGAE filaments that may arise from stacking of tyrosine residues ^27^.

Structural variation between DSB(+) and DSB(-) filaments was further investigated by examining surface exposure, using protease resistance to Proteinase K (PK) digestion ^28^. Fibrils formed from dGAE, dGAE+DTT and dGAE-C322A at 400μM were diluted to 200μM and incubated with 25μg/ml PK for 1h at 37°C and the products examined using SDS-PAGE (Figure 3ci). Without PK digestion, there is a noticeable difference in the SDS soluble species between conditions. dGAE filaments are SDS soluble and mostly run as apparent dimers, shown at 20kDa and 24kDa, respective to their 10-kDa and 12-kDa monomers shown in our previous work ^19^. We also see minor band at a higher molecular weight band around 40kDa and another lower at ∼17kDa that is not seen for the dGAE+DTT or dGAE-C322A fibril samples. dGAE+DTT and dGAE-C322A runs with a far less intense band at 24kDa and a stronger band at 12kDa, representing monomeric dGAE (green arrow). The very weak 24-kDa band present in the dGAE+DTT and dGAE-C322A samples must be disulphide-independent dimers. Post PK digestion, dGAE filaments formed in non-reducing conditions run at 8kDa (red arrow), showing a truncated dGAE monomer consisting of the protease-resistant core, and another band that is slightly higher molecular weight than 12kDa, this could represent the full-length dGAE monomer (12kDa). Mass spectrometry of the 8-kDa band showed that this fragment maps to a protease-resistant core of dGAE filaments encompassing the region His299 – Lys370 (Sup Figure 3a). These two distinct bands post-PK treatment were observed even with an increase in PK concentration up to 250µg/ml, suggesting that their presence of was not due to incomplete digestion (Sup. Figure 3b). dGAE+DTT and dGAE-C322A fibrils run with a sole band at 8kDa and showed a significantly more when compared to the dGAE fibrils (Figure 3c ii). Overall, the PK digestion data highlight differences in the structural core of the fibrils formed from DSB(-) and DSB(+) dGAE fibrils.

These observations may suggest that the dGAE sample has two populations of filaments, one with DSB and one without DBS. The latter population would show filaments having similar characteristics to the those formed in the dGAE+DTT and dGAE-C322A samples, where DSB is inhibited. The filaments can be partially digested by PK before being broken down into monomers by SDS, which yields the band at 8kDa. The other population consists of a filament structure that does not undergo complete digestion of dGAE by PK but is affected sufficiently by the PK for it to become more SDS soluble. This is because there is no 12-kDa band without PK, but a strong 12-kDa band with PK. Quantification of the 12-kDa vs 8-kDa band of dGAE fibrils with PK showed us that there is generally more of the 12-kDa band than the 8-kDa band representing the core region seen in the fibrils formed without DSB. This difference was not significant but demonstrates that there is a greater presence of the fibrils giving rise to 12-kDa band, than those responsible for the 8-kDa band (Figure 3c iii).

These data suggest that DSB has a significant effect on the macromolecular structure of dGAE filaments that could explain the differences in assembly kinetics and seeding propensity observed previously. DSB(-) fibrils show a distinct CD spectrum, SDS-PAGE profile, and susceptibility to protease digestion when compared to DSB(+) fibrils.

### DSB(-) species can recruit endogenous tau in FRET biosensor cells

Having shown that DSB(-) dGAE species are more capable of seeding assembly compared to DSB(+) dGAE species *in vitro*, we investigated whether dGAE aggregates are able to recruit endogenous tau within cells. We utilised the Tau RD P301S FRET Biosensor model of tau aggregation, in which HEK293T cells express two populations of tau corresponding to the repeat domain (RD) region and carrying the P301S mutation associated with FTD ^24^. Each population of tau has a separate fluorescent tag, either CFP or YFP. The addition of seed-competent tau leads to the aggregation of endogenous tau and the resultant close proximity of the two populations of tau results in a FRET signal, which can be interpreted as the aggregation of endogenous tau.

We utilised the automated ImageXpress Pico system to develop a highly sensitive method of measuring and analysing the FRET signal in live FRET Biosensor cells (Sup. Figure 5). Cells were plated in a 96-well plate and treated with soluble dGAE or dGAE-C322A, and sonicated or non-sonicated fibrils (Figures 2 and 3, respectively). The addition of non-aggregated soluble dGAE protein resulted in no appreciable FRET signal intensity, suggesting that soluble dGAE and dGAE-C322A did not recruit endogenous tau to assemble during the time-course of the experiment (Figure 4a and b i). DSB(+) dGAE fibrils and sonicated fibrils did not induce a FRET signal, suggesting that the DSB(+)species were unable to recruit endogenous tau. However, there was a significant increase in the FRET signalling with the incubation of dGAE+DTT and dGAE-C322A fibrils (DSB (-)) and sonicated fibrils when compared to the DSB(+) dGAE fibrils and sonicated fibrils (Figure 4a and b). This suggests that only DSB(-) fibrils can recruit cellular endogenous tau to form aggregates.

**Figure 4:**
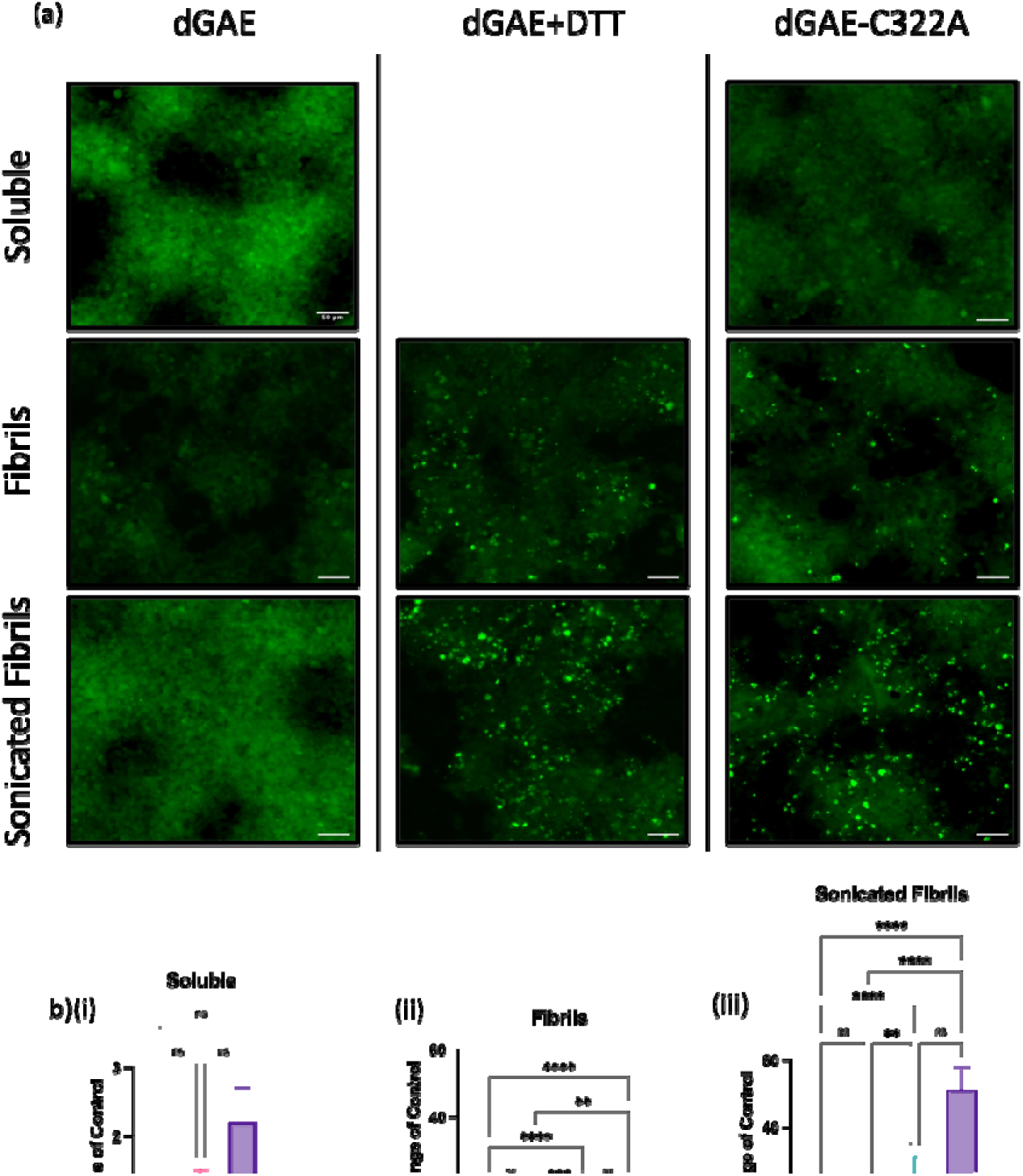
dGAE+DTT and dGAE-C322A induce signal in FRET biosensor cells. FRET Biosensor cells plated in 96-well plate and treated extracellularly with 10μM of each of the dGAE species for 3d before imaging. (a) Images of FRET Biosensor cells taken from each condition using an ImageXpress Pico. The FRET signal appears as a punctate green fluorescence and diffuse green signal seen in cells without FRET signal. Scale bars represent 50μM. Data not shown for soluble dGAE-DTT. (b) Quantification and comparison from analysis of FRET signal in response to soluble dGAE and dGAE-C322A when compared to control. Kruskal-Wallis test showed no significant difference between samples (p = 0.1396). (c) Quantification and comparison from analysis of FRET signal induced by dGAE, dGAE+DTT and dGAE-C322A fibrils when compared to control. Kruskal-Wallis test showed no significant difference between samples (p < 0.0001). Dunn’s Multiple comparison test showed dGAE-C322A fibrils (9.825 ± 1.60) induced a significant increase in FRET signal when compared to control (1.00 ± 0.07265, p < 0.0001) and dGAE fibrils (1.795 ± 0.2744, p = 0.0012). dGAE+DTT (11.19 ± 1.917) fibrils induce a significant increase in FRET signal compared to control (p < 0.0001) and dGAE fibrils (p = 0.0003). No significant difference between dGAE+DTT and dGAE-C322A fibrils. (d) Quantification and comparison from analysis of FRET signal induced by dGAE, dGAE+DTT and dGAE-C322A sonicated fibrils when compared to control. Kruskal-Wallis test showed no significant difference between samples (p < 0.0001). Dunn’s multiple comparison test showed dGAE-C322A fibrils (51.15 ± 6.726) induced a significant increase in FRET signal when compared to control (1.00 ± 0.07265, p < 0.0001) and dGAE fibril (2.689 ± 0.2828, p < 0.0001). dGAE+DTT (25.86 ± 5.554) fibrils induced a significant increase in FRET signal compared to control (p < 0.0001) and dGAE fibrils (p = 0.0092). There was no significant difference between dGAE+DTT and dGAE-C322A fibrils.

The above findings provide further evidence for distinct differences in the macromolecular structure of fibrils formed in the absence of DSBs that facilitate their ability to seed aggregation. The addition of sonicated fibrils caused a significant increase in the FRET signal compared to non-sonicated fibrils, potentially because of their smaller size making it easier for their internalisation within the cell.

## DISCUSSION

In this study, we aimed to evaluate the self-assembly mechanisms involved in dGAE amyloid formation and the contributions of DSB to these mechanisms, filament structure and role in tau pathology. We first optimised conditions for reproducible self-assembly of the dGAE tau fragment in environments that either favour DSB or inhibit DSB (using reducing conditions with 10mM DTT and a cysteine variant of dGAE). ThS fluorescence assays from a monomeric solution showed that dGAE aggregation demonstrated the expected sigmoidal curve with a pronounced lag time followed by a sudden elongation phase and a reaction order close to -0.5, which suggest that secondary processes (secondary nucleation and fragmentation) dominate the assembly process. The dGAE aggregation kinetics fitted closely with the aggregation models of multi-step secondary nucleation and saturating elongation with fragmentation (Figure 1) and the seeding assays showed a clear accelerated assembly with the addition of aggregates (Figure 2). The reduction, but not the elimination, of the lag phase is evidence for a surface-mediated seeding process (multi-step nucleation). Surface-mediated assembly is still a nucleation-dependent process, requiring a slow nucleation phase, albeit enhanced reaction when compared with the absence of seeds. This has been reported for Aβ42 monomer and seeds produced from the yeast prion-forming protein, Sup35NM ^26^. Experimentally, we also show that sonicated fibrils were more competent seeds when compared to fibrils (Sup. Figure 3). We predicted that a seed containing larger fibril surface, such as long fibrils, should facilitate the multi-step secondary nucleation process that occurs on the fibril surface, whereas a seed with more exposed ends, such as sonicated fibrils, should facilitate the elongation with fragmentation process that occurs at the ends of fibrils. This suggests that the sonicated fibrils, with more exposed ends, favoured the elongation and fragmentation processes, suggesting the presence of fragmentation in dGAE aggregation. However, it is difficult to determine whether the sonication aided seeding due to number of seeds or aiding their availability by influencing their size. Nevertheless, it is clear that sonication has a significant effect on increasing the seeding ability of dGAE-C322A seeds *in vitro*, which is later also shown in FRET Biosensor cells. It is also important to consider the involvement of agitation in our experimental conditions since we were unable to obtain reproducible spontaneous assembly of dGAE without agitation. The agitation of the sample to induce reproducible aggregation kinetics may cause fragmentation of aggregated species ^29^. Considering all the evidence we have gathered for dGAE aggregation in the absence and presence of seeds, we suggest that both elongation with fragmentation and multi-step secondary nucleation are active in the process of dGAE aggregation. These processes are not mutually exclusive, and it is important to highlight their presence within a complex assembly process that makes it difficult to draw a conclusion on the precise mechanism involved.

Investigating the self-assembly of amyloidogenic proteins under varying conditions helps to highlight important mechanisms during the assembly process, and how that translates to the formation of pathological aggregates found in the brain. DSB has been a debated topic for tau assembly, with studies using previous models of *in vitro* tau aggregation (T40 and K18/K19) showing DSB being essential for self-assembly ^17^, whereas our previous work using dGAE has shown that inhibition of DSB enhances self-assembly ^19^. This study has provided a more detailed mechanistic investigation, using ThS aggregation traces, to study the impact of inhibiting DSB. Our results show that the inclusion or inhibition of DSB has little effect on the global mechanism of assembly for dGAE, with secondary nucleation processes dominating the assembly reaction (Figure 1 and Sup. Figure 2). However, there is a clear acceleration of aggregation through a significant reduction in the lag phase when DSB is inhibited. This suggests that DSB does not affect the overall mechanism of self-assembly, but the absence of DSB enhances the aggregation of dGAE and reduces the lag phase of assembly. This led us to investigate the ability of the species formed from each condition to act as a seed. We found that DSB(-) dGAE seeds are more seed competent than DSB(+) species, in both *in vitro* aggregation (Figure 2) and in recruiting endogenous tau in the FRET biosensor cells (Figure 4). This suggests that there are subtle differences in the structure of the fibrils produced under different conditions that results in a species with varying seeding capabilities to facilitate the secondary processes of assembly observed with our mechanistic studies. With these observations, we have devised a model to explain the enhanced assembly observed in the dGAE+DTT and dGAE-C322A samples and the increased ability of DSB(-) species at seeding aggregation (Figure 5). We propose that the inhibition of DSB during dGAE assembly produces dGAE aggregates that are more seed-competent, which accelerates further aggregation. Whereas, allowing the formation of DSB results in far fewer of these seed-competent species, which shows in a much slower assembly reaction. This is also supported with the finding that secondary processed dominant within the assembly reaction whether DSB is allowed or inhibited, meaning that these seed-competent species play a crucial role in dGAE assembly. PK digestion assays in figure 3 also suggested that there are two populations of filaments with different proteolytic profiles, comprising of the DBS(+) and DBS(-) species, which aren’t observed in the sample inhibiting DSB.

**Figure 5:**
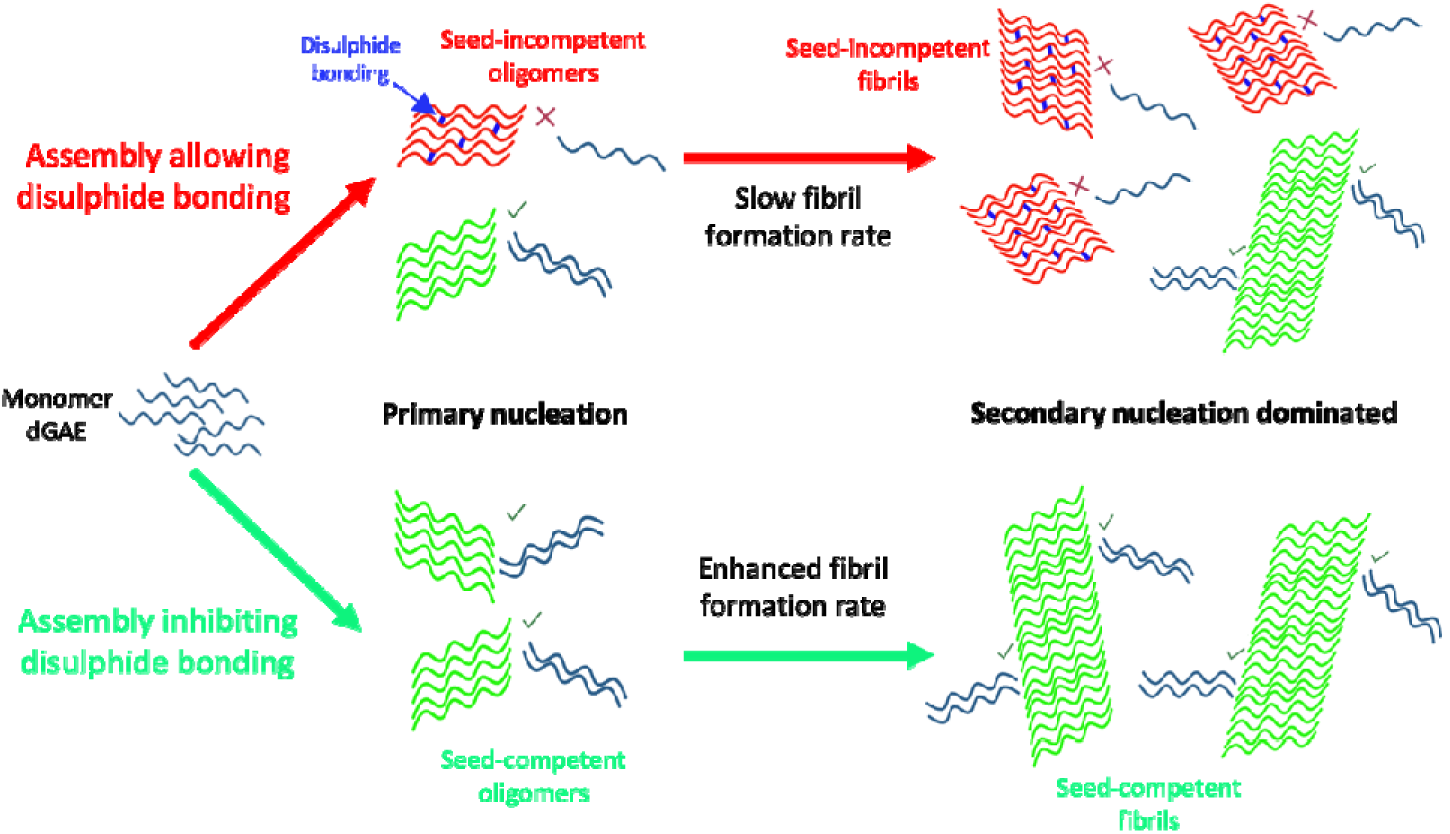
Schematic representation of the assembly of dGAE in non-reducing and reducing conditions throughout the assembly process and the effect of disulphide bonding on these processes. We speculate that the assembly in non-reducing conditions results in a mixture of oligomers after primary nucleation with and without disulphide bonding. The oligomers with disulphide bonds (illustrated in red with blue connections) are less seed incompetent, whilst the oligomers without disulphide bonds are more seed competent (shown in green). Due to less seed-competent oligomers, the secondary dominant process into mature fibrils is slow, resulting in a mixture of seed-incompetent and seed-competent fibrils. In DSB(-) conditions, disulphide bonds are inhibited which leads to the primary nucleation of seed-competent oligomers only. This means that secondary nucleation is facilitated and there is an enhanced fibril formation. This results in a sample of fully seed-competent fibrils. Created with BioRender.com.

Further differences in the structure of the fibrils were observed using TEM, CD and SDS-PAGE and proteolysis analysis (Figure 3). CD results show two distinct spectra between the fibril types, with the fibrils formed in the absence of DSB containing two additional peaks at ∼250nm and ∼290nm, which may be due to stacking of tyrosine residues ^27^. SDS-PAGE illustrated differences in the presence of species in the DSB(-) and DSB(+) aggregates post-SDS degradation. A strong band at 24kDa (indicating dGAE dimer) ^19^ is present with dGAE fibril sample, whereas the dGAE+DTT and dGAE-C322A fibrils have a strong 12-kDa band (indicative of dGAE monomer) together with a weaker dimer band. Previously, we reported that dGAE fibrils formed under reducing and non-reducing conditions differed in terms of the extent of core using cross polarisation and INEPT solid-state NMR ^30^. Fibrils formed without DTT were more dynamic suggesting that there was less of the protein incorporated into a stable core compared with the well-ordered, static core found in dGAE+DTT fibrils. In this study, PK treatment was used to further investigate differences in structure and protease resistance of the fibrils. dGAE+DTT and dGAE-C322A fibrils exhibit one strong band at 8kDa post-treatment. This is likely to be the partially digested dGAE monomer that forms a protease-resistant, but SDS-soluble, core of the fibrils formed in the absence of DSB. However, in the dGAE filaments, we see the presence of two distinct bands. One band at 8kDa, representing the protease-resistant core, and the other at 12kDa, representing undigested dGAE monomer. At first, this was assumed to be due to insufficient PK resulting in undigested protein. But the presence of these two bands persisted for PK concentrations up to 250µg/ml suggesting that this is not the case (Sup. Figure 4). We propose that the dGAE sample contains two separate populations of fibrils. One fibril population is formed without DSB, which is the dominant fibril seen in the dGAE+DTT and dGAE-C322A sample and is found as an 8-kDa band following PK digestion. The other fibril population is formed with DSB and has a conformation that is resistant to PK. Following electrophoresis, undigested dGAE monomer at 12kDa is present, but PK is still able to affect their conformation by making the filaments more SDS soluble, shown by a lack of dimer post-PK treatment (Figure 6).

**Figure 6:**
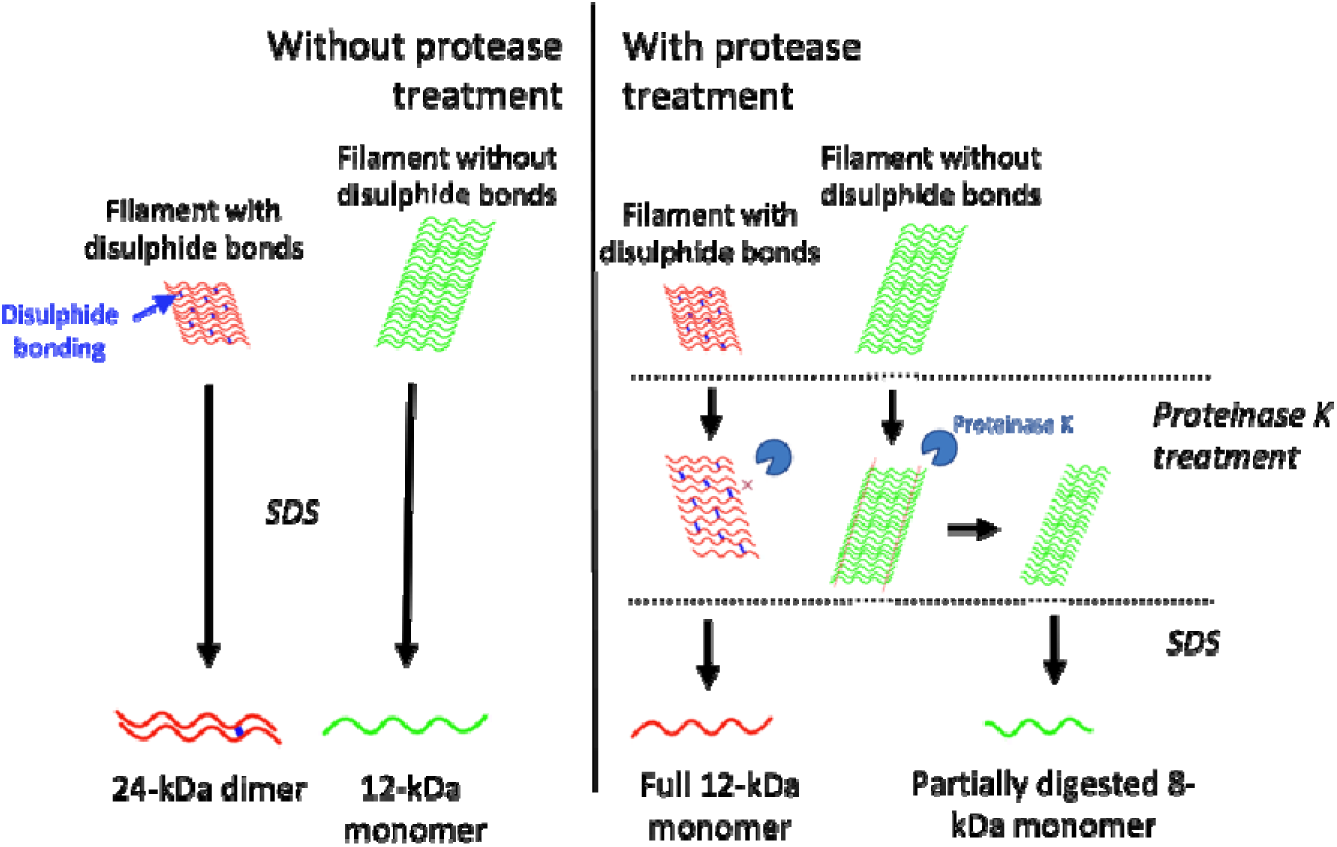
Illustrating the proteinase-K and SDS solubility of filaments formed with and without disulphide bonds as observed with SDS-PAGE. Filaments with disulphide bonds (red filaments with blue connections) are broken down mainly to dGAE dimers (24kDa), seen with SDS-PAGE, whereas filaments without disulphide bonds (green filaments) are broken down to dGAE monomers (12kDa) after separation by SDS-PAGE. Proteinase K activity is not able to partially digest filaments with disulphide bonds, but it does make them more SDS soluble, producing a dGAE monomer (12kDa) with SDS-PAGE. In contrast, filaments without disulphide bonds are less resistant to proteinase K activity, resulting in partially digested monomer, which are then seen as 8-kDa monomers with SDS-PAGE. Created with BioRender.com

To gain further insights into the relevance of the seeding process for tau pathology, we utilised FRET biosensor cells to investigate which species were able to seed the assembly of endogenously expressed forms of tau. Results revealed that only fibrillar, and not monomeric, forms were able to seed assembly and that sonicated fibrils were more efficient at seeding than non-sonicated fibrils. Furthermore, it appeared that DSB(+) dGAE species were unable to seed endogenous tau assembly, whether fibrillar or sonicated. In contrast, DSB(-) dGAE species were effective in the recruitment of endogenous tau in the FRET Biosensor cells without the need for agents, such as Lipofectamine® 2000, to aid internalisation. Sonicating the fibrils resulted in a significant increase in FRET signalling, suggesting the sonicated fibrils are superior at seeding the endogenous tau. This could be due to the smaller size of the seeds are better at internalising within the cell, or that the smaller dGAE species are better at seeding, which was observed in vitro (Sup. Figure 3). These results indicate that dGAE aggregates are able to enter cells and recruit endogenous tau, further highlighting the use of dGAE aggregates for the study of tau seeding and propagation. The various profiles of species present after PK digestion might also suggest that DSB(-) dGAE aggregates are more protease resistant and, therefore, more likely to persist within the cell to recruit endogenous tau. This is consistent with a proteolytic selection process discussed by Bansal and colleagues, who state that fibril formation results in mixture of polymorphic filaments that vary in their protease stability, and proteolytic activity leads to the degradation of soluble proteins resulting in only protease-stable filaments ^31^. Our data suggest that tau fibrils formed with DSB could be more susceptible to digestion, whereas the DSB(-) fibrils are more protease-resistant and able to persist in the cell. These fibrils also exhibit the pathological characteristics of being more seed-competent which we might expect facilitate the recruitment of endogenous tau for the propagation of tau pathology throughout the brain.

Taken together, our data suggest that inhibition of DSB is a key step in the self-assembly of dGAE and the production of pathological seed-competent species of tau. Identifying the inhibition of DSB as an essential pathological step in dGAE aggregation is consistent with studies investigating tau aggregates in disease. Cryo-EM structures of PHFs in AD revealed that the cysteine residues are buried deep within the structure of the filament and therefore unavailable for DSB, which strongly indicates its lack of relevance in disease filament formation ^5^. In addition, the predominantly reducing environment of the cell cytosol ^32^ would favour the inhibition of DSB and the formation of these more seed-competent species that we have shown with dGAE+DTT and dGAE-C322A. We propose that the balance between the structures of the fibrils formed through DSB or by inhibiting DSB is responsible for the speed of aggregation of dGAE and the seeding capabilities of the aggregated dGAE species in vitro and in FRET Biosensor cells.

In conclusion, we have utilised the dGAE fragment to investigate the role of DSB in tau self-assembly. We suggest that the inhibition of DSB is an important step towards pathological self-assembly of dGAE and the formation of aggregated tau species capable of propagating tau pathology. Our data indicate that this is due to a distinct difference in fibril structure as a result of the inhibition of DSB, which produces a more seed-competent fibril shown within in vitro assembly assays and within a tau aggregation cellular model. Recent cryo-EM studies have illustrated that subtle, but distinct, changes in the conformation of tau fibrils throughout tauopathies, highlighting the importance of structure in the study of tau pathology. This has now been extended to the dGAE tau aggregation model, illustrating that varying the assembly conditions can have a drastic effect on the assembly and fibril characteristics. This further demonstrates the use of the dGAE in vitro model in investigating the mechanisms that play a pivotal role in the stages of aggregation and tau propagation and in the development of tau-targeted therapies.

## ASSOCIATED CONTENT

### Supporting Information

The Supporting Information is available free of charge on the ACS Publications website.

Enclosed in .pdf

Figure S1.Fitting of global assembly mechanisms to ThS kinetics traces.

Figure S2. Primary and secondary rate constants of assembly from each condition.

Figure S3. The effect of sonication on seeding ability of dGAE-C322A seeds.

Figure S4. The basis for fibrillar Protease K resistance.

Figure S5. Raw images and analysis overlay of FRET Biosensor cells acquired using ImageXpress Pico and analysed in CellReporterXpress

## AUTHOR INFORMATION

### Present Addresses

†If an author’s address is different than the one given in the affiliation line, this information may be included here.

### Author Contributions

The manuscript was written through contributions of all authors. All authors have given approval to the final version of the manuscript.

## ACKNOWLEDGMENT

We thank Dr Pascale Schellenberger for support with electron microscopy. We would also like to thank Kate Heesom from the Proteomics facility at the University of Bristol for her mass spectrometry work.

## ABBREVIATIONS

(YT): Yeast Tryptone 2x
(IPTG): Isopropyl β-D-1-thiogalactopyranoside
(EDTA): Ethylenediaminetetraacetic acid
(DTT): 1mM dithiothreitol
(EGTA): ethylene glycol-bis(2-aminoethylether)-N,N,N’,N’-tetraacetic acid
(PB): phosphate buffer
(ThS): Thioflavin S
(BCA): Bicinchoninic acid
(TEM): Transmission Electron Microscopy
(CD): Circular dichroism
(PK): proteinase K
(SDS-PAGE): Sodium dodecyl-polyacrylamide gel electrophoresis

