## Supplemental info for "Inhibiting disulphide bonding in truncated tau297-391 results in enhanced self-assembly of tau into seed-competent assemblies"

Supplementary Figure 1.

Nucleation Elongation

Secondary Nucleation

Saturating Elongation and Secondary Nucleation

dGAE

dGAE+DTT

dGAE-C322A

a)(i)

b)(i)

c)(i)

(ii)

(ii)

(ii)

(iii)

(iii)

(iii)

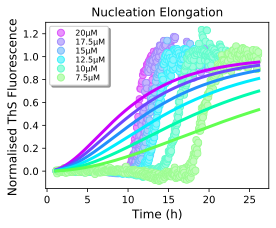

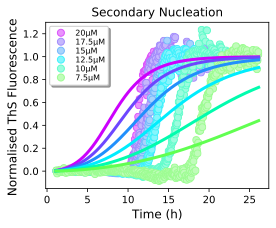

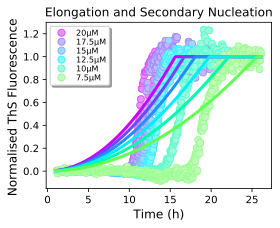

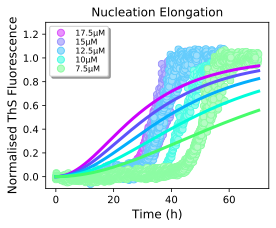

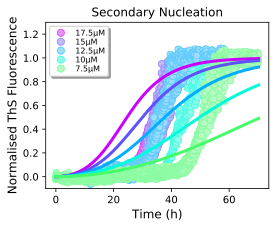

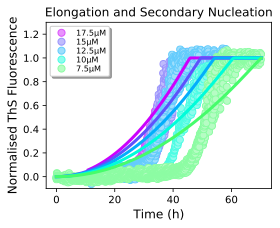

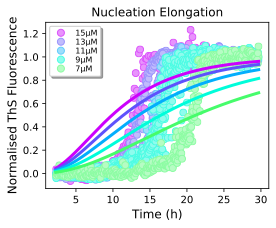

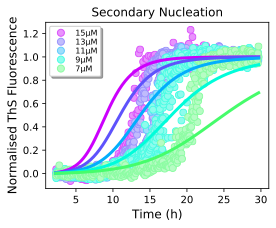

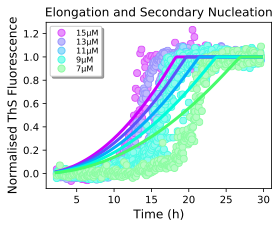

**Supplementary figure 1: Global assembly mechanisms that do not fit dGAE aggregation kinetics.**

**dGAE assembled in non-reduced conditions shown on left section.** (i), dGAE assembled in reduced conditions with 10mM DTT in middle section (ii), and dGAE-C322A assembled in non-reduced conditions (iii). Normalised kinetic profiles plotted against different models of assembly:(a) nucleation elongation, (b) secondary nucleation, and (c) saturating elongation and secondary nucleation

Supplementary figure 2:

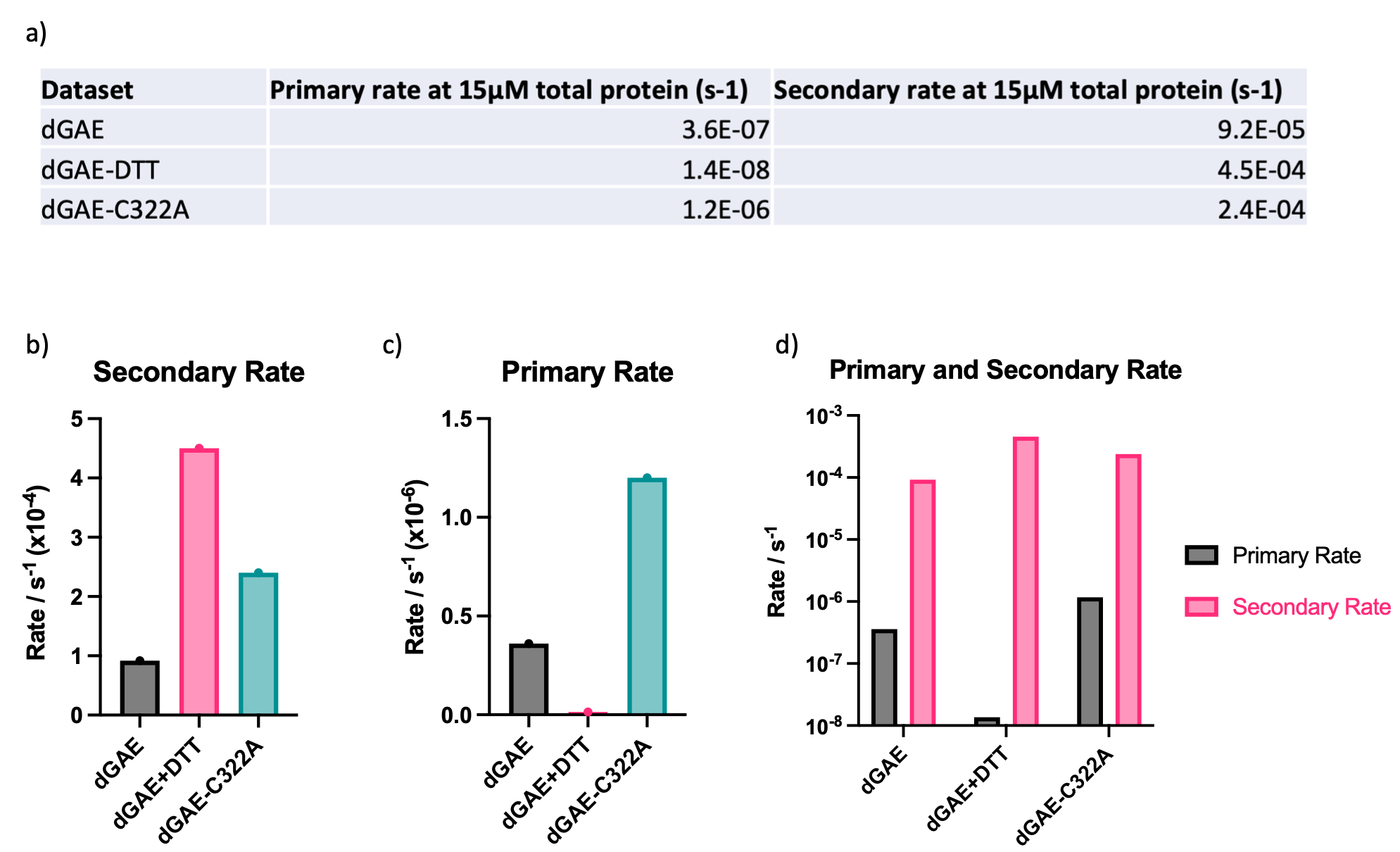

**Supplementary Figure 2: The primary and secondary rate constants were obtained from the global fit data shown in figure 1 using Amylofit software using the traces of 15µM experiments**. (a). The secondary rate (b) and primary rate (c) are shown separately to illustrate the differences between the assembly conditions and shown all together for comparison between primary and secondary within each assembly conditions.

Supplementary Figure 3

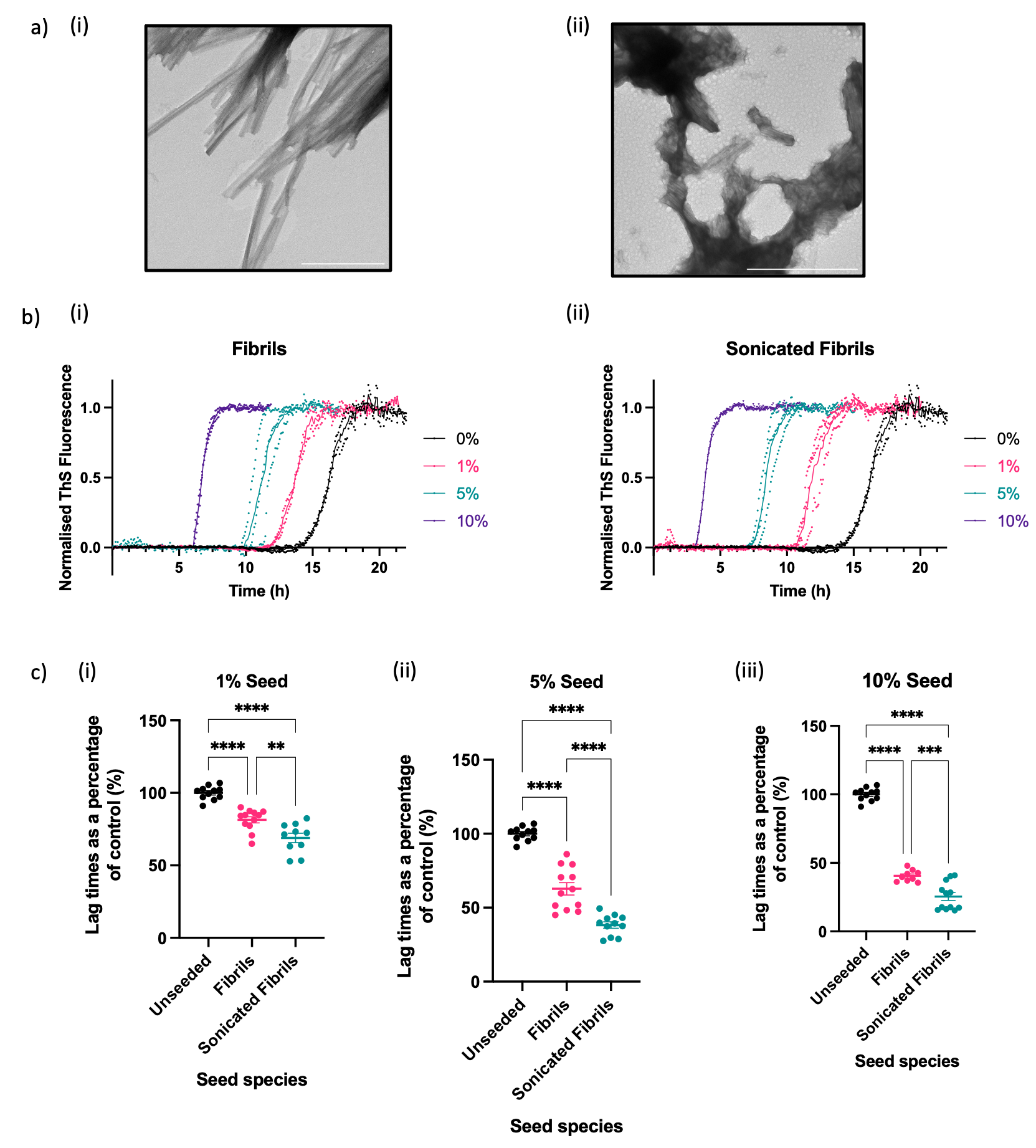

**Supplementary Figure 3: Seeding from dGAE-C322A seeds before and after sonication**

Electron micrograph of dGAE-C322A fibrils produced with 400μM monomeric dGAE-C322A (ai) and after sonication (aii). Scale bar represents 500nm. Example of normalised thioflavin-S kinetics from a single experiment showing the seeding capability of dGAE-C322A fibrils (bi) and after sonication (bii). 0% (control - black), 1% (pink), 5% (green) and 10% (purple). (ci) Quantification and comparison of the lag times with 1% seeds of each condition as a percentage of the control. One-way ANOVA shows significant difference between groups (F=43.80, R^2^=0.7449, p < 0.0001) from 3 independent tests. Turkey’s multiple comparison test shows sonicated fibril seed (68.92% ± 3.247%) induce a significant reduction in lag phase when compared to the control (100% ± 1.399%, p < 0.0001) and fibril seeds (81.46% ± 2.142%, p = 0.0018). Fibril seeds induce a significant reduction in lag phase when compared to control (p < 0.0001). (cii) Quantification and comparison of the lag times with 5% seeds of each condition as a percentage of the control. One-way ANOVA shows significant difference between groups (F=110.9, R^2^=0.8774, p < 0.0001) from 3 independent tests. Turkey’s multiple comparison test shows sonicated fibril seed (38.25% ± 2.127%) induce a significant reduction in lag phase when compared to the control (100% ± 1.399%, p < 0.0001) and fibril seeds (62.76% ± 4.140%, p < 0.0001). Fibril seeds induce a significant reduction in lag phase when compared to control (p < 0.0001). (ciii) Quantification and comparison of the lag times with 10% seeds of each condition as a percentage of the control. One-way ANOVA shows significant difference between groups (F=338.6, R^2^=0.9589, p < 0.0001) from 3 independent tests. Turkey’s multiple comparison test shows sonicated fibril seed (25.47% ± 2.920%) induce a significant reduction in lag phase when compared to the control (100% ± 1.399%, p < 0.0001) and fibril seeds (40.51% ± 1.403%, p = 0.0002). Fibril seeds induce a significant reduction in lag phase when compared to control (p < 0.0001).

Supplementary Figure 4.

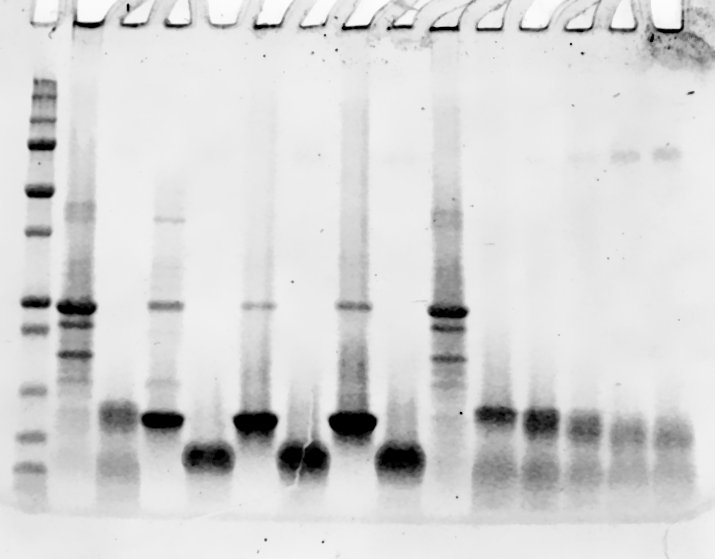

10

5

15

20

25

37

50

PK Conc.

(μg/ml)

0

10

25

50

100

250

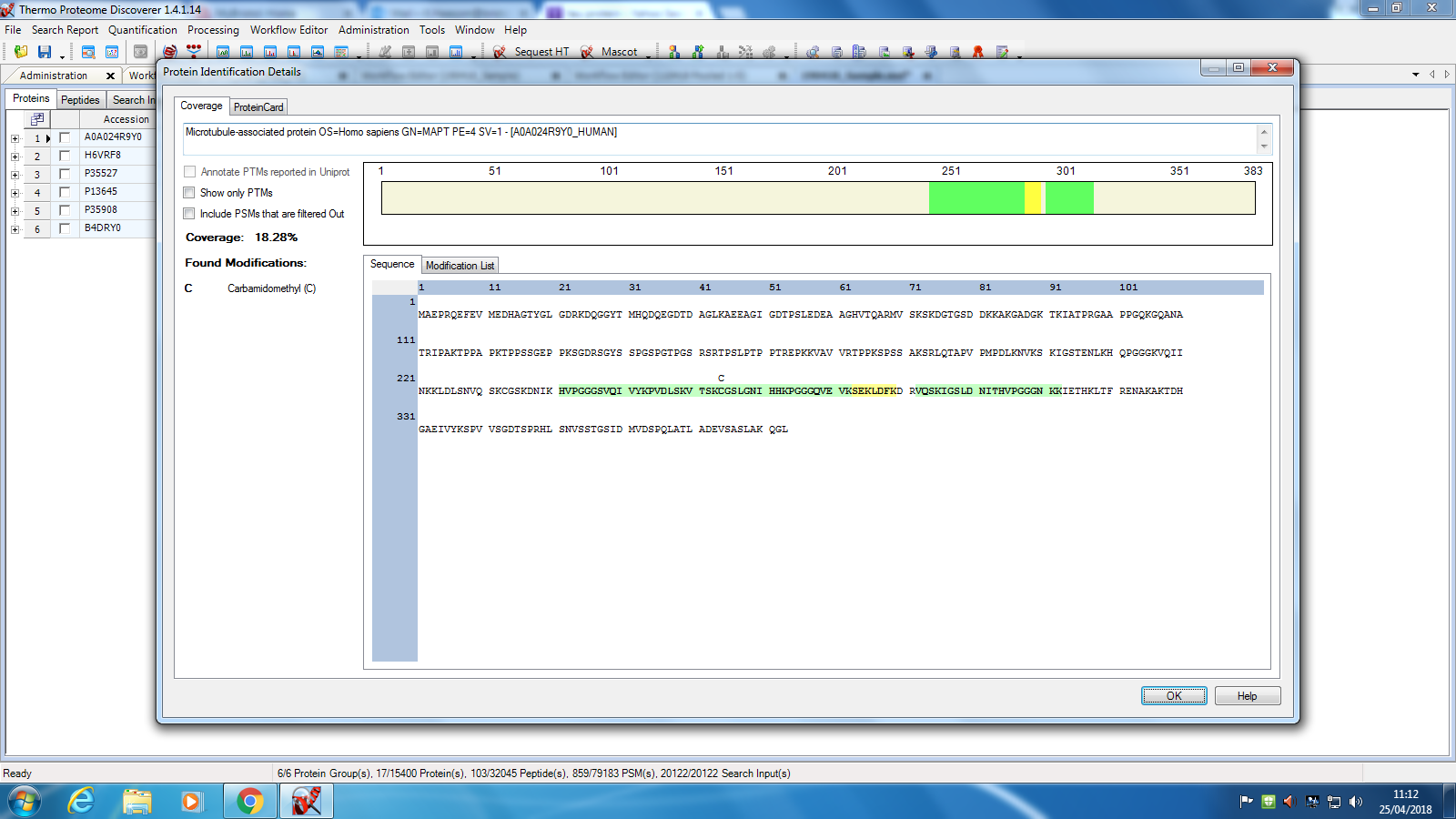

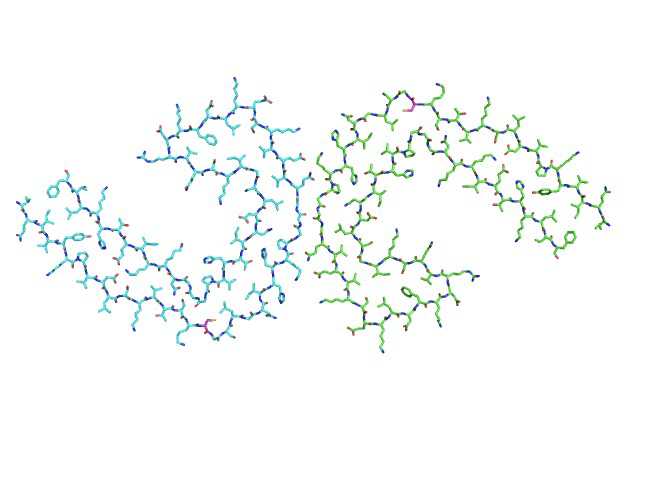

I

Q

G

G

G

P

H

K

I

306

311

Cys322

297

337

H299

K370

a)(i)

(ii)

b)

**Supplementary figure 4: 12kDa band is resistant to increased protease K concentration.**

(ai) Mass spectrometry analysis of the band at 8kDa after protease K treatment. Green region shows the region believed to be protease K resistant in comparison with the whole dGAE sequence indicated with the blue box. (aii) Illustrating the protease K resistant core using the dGAE fold resolved by Lövestam et al., 2022. (b) 200μM dGAE fibrils formed in non-reducing conditions incubated with 0, 10, 25, 50, 100 and 250μg/ml protease K for 1h at 37℃ before being analysis with SDS-PAGE and stained with Coomassie protein stain.

Method

Mass spectrometry carried out to identify the sequence of the 8 kDa band after PK digestion. The band was cut out of the gel and put into water in a microcentrifuge tube. The excised band was sent for tryptic digestion and LC-MS analysis by the Proteomics facility at the University of Bristol [1].

Supplementary Figure 5.

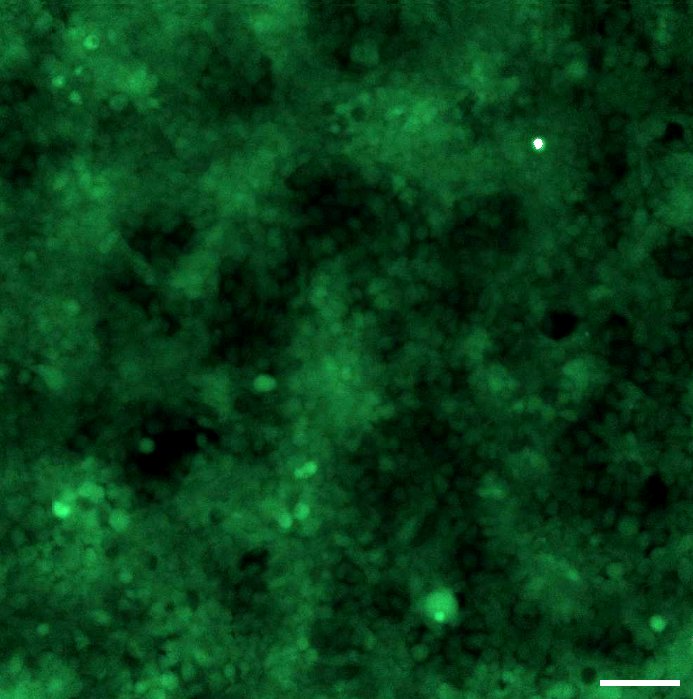

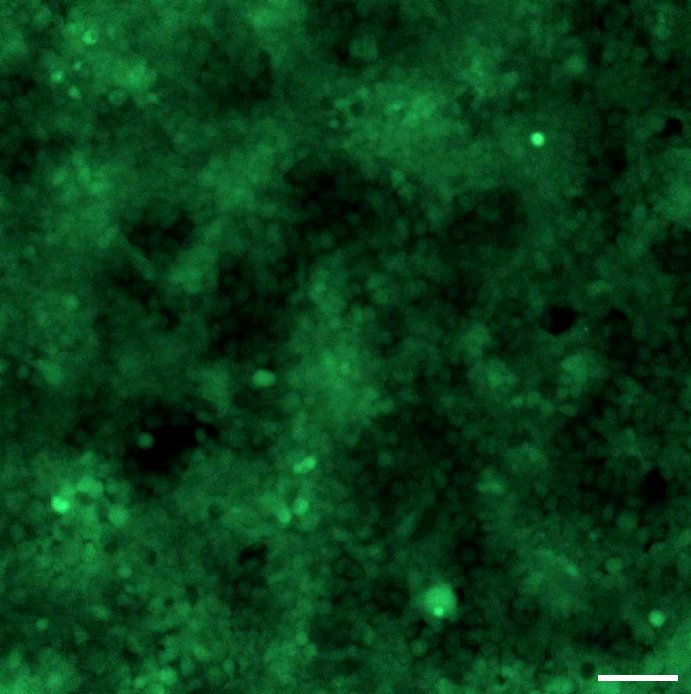

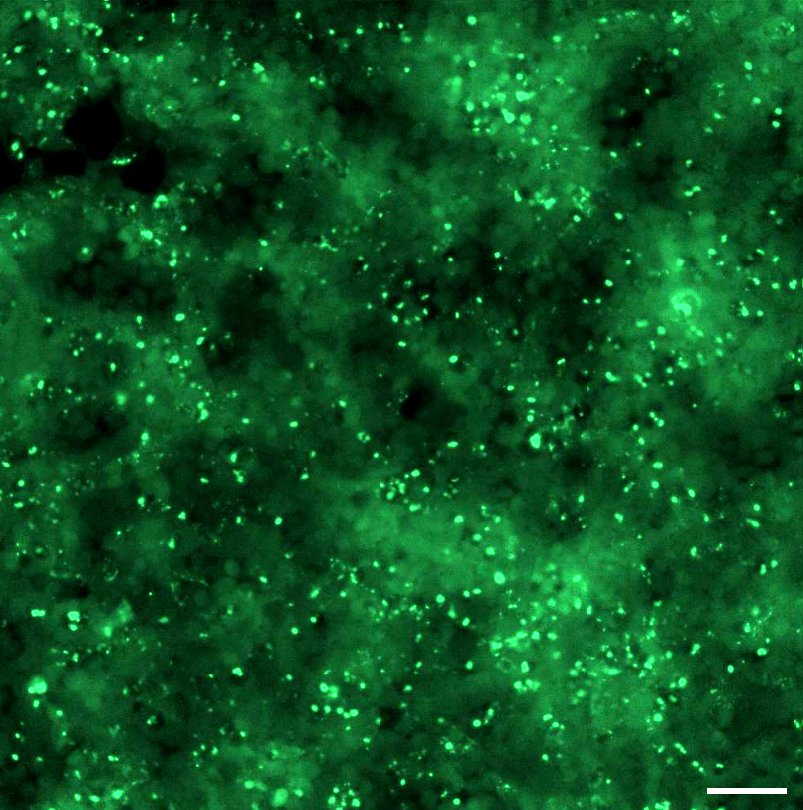

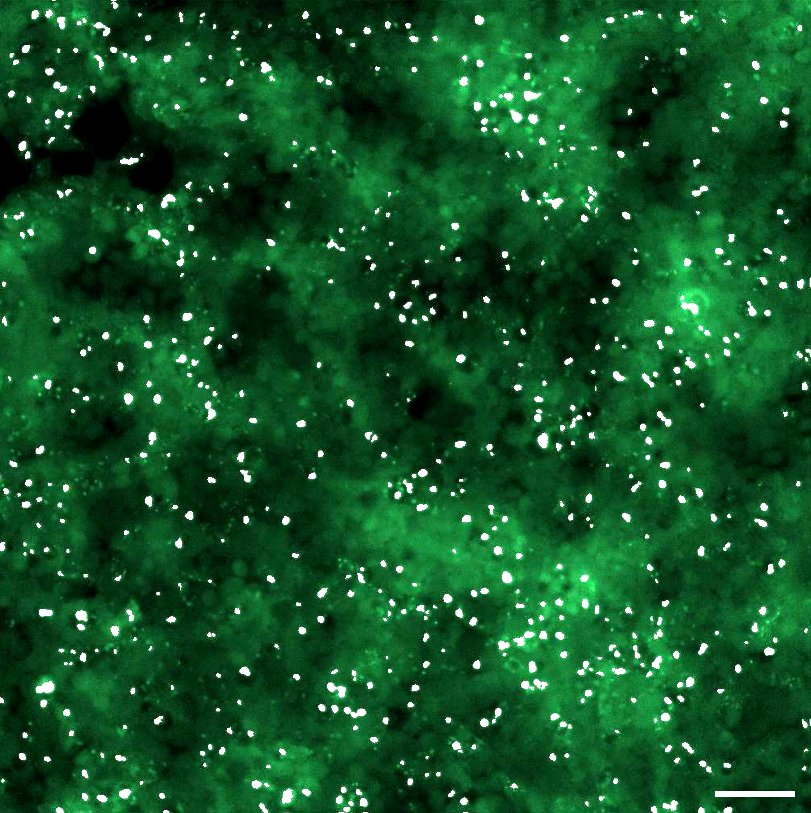

Control

Seed

Raw image

Analysis overlay

**Supplementary Figure 5: Demonstration of image analysis undertaken of FRET Biosensor cells with the Molecular Devices ImageXpress Pico and CellReporterXpress.**

Raw images of FRET Biosensor cells treated with phosphate buffer along (control) or with 10μM sonicated dGAE-C322A fibrils (seed), which shows an increase in FRET signal taken with the ImageXpress Pico without the analysis overlay (top row). Images acquired with the analysis overlay undertaken (white signal shows isolated FRET signal) with the described parameters to isolate the FRET signal wanted. All scale bars represent 50μm.
